# Anisotropy Measure from Three Diffusion-Encoding Gradient Directions

**DOI:** 10.1101/2021.03.24.436786

**Authors:** Santiago Aja-Fernández, Guillem París, Antonio Tristán-Vega

## Abstract

**Purpose:** We propose a method that can provide information about the anisotropy and orientation of diffusion in the brain from only 3 orthogonal gradient directions without imposing additional assumptions.

**Methods:** The method is based on the Diffusion Anisotropy (DiA) that measures the distance from a diffusion signal to its isotropic equivalent. The original formulation based on a Spherical Harmonics basis allows to go down to only 3 orthogonal directions in order to estimate the measure. In addition, an alternative simplification and a color-coding representation are also proposed.

**Results:** Acquisitions from a publicly available database are used to test the viability of the proposal. The DiA succeeded in providing anisotropy information from the white matter using only 3 diffusion-encoding directions. The price to pay for such reduced acquisition is an increment in the variability of the data and a subestimation of the metric.

**Conclusions:** The calculation of anisotropy information from DMRI is feasible using fewer than 6 gradient directions by using DiA. The method is totally compatible with existing acquisition protocols and it may provide complementary information about orientation in fast diffusion acquisitions.

## 1 INTRODUCTION

The term Diffusion Magnetic Resonance Imaging (DMRI) refers to a set of diverse imaging techniques that, applied to brain studies, provide useful information about the organization and connectivity of the white matter. Although the most relevant feature of DMRI is its ability to measure orientational variance in the different tissues, i.e. anisotropy, it is still a complementary practice to some structural studies to perform a fast acquisition to obtain a measure of the *amount* of diffusion. A common implementation in commercial scanners, like EPI-DWI, acquires only 3 separate orthogonal diffusion weighted images (DWIs) with diffusion gradients aligned with directions (*x, y, z*). These 3 DWIs are averaged into a final combined image [1] that resembles measures like the Mean Diffusivity (MD) [2] or the Average Sample Diffusion [5]. Due to the limitation in the number of gradient directions, no extra information is provided. If a measure of anisotropy and orientation of the diffusion wants to be extracted, there is a minimum requirement of 6 acquired DWIs in order to estimate the components of the diffusion tensor (DT) [3]. Under the DT approach, it would still be possible to calculate an anisotropy measure with fewer than 6 gradient directions, but we must impose a restricted model that reduces the number of values to estimate, like, for instance, assuming that the diffusion has a cylindrical symmetry.

In this paper we propose a new method that can provide information about the anisotropy in the diffusion from only 3 orthogonal gradient directions without imposing additional assumptions to the DT model. This method is totally compatible with existing fast diffusion acquisition, since it only makes use of the same 3 DWIs already acquired. This way, no extra scanning time is needed: the same sequence that provides MD images can also provide anisotropy information.

The method is based on a novel anisotropy metric called Diffusion Anisotropy (DiA) proposed in [4]. The metric measures the distance from the actual diffusion signal to its isotropic equivalent. Its original formulation relies on the fitting on the signal using a basis of Spherical Harmonics (SH), but an alternative simpler formulation is here proposed to be exclusively used with 3 orthogonal gradient directions. In addition, we also present a color-coding method, similar to the one used for the Fractional Anisotropy (FA) in DT imaging. We carry out some examples and tests to show that, although the variability of the anisotropy image is high (compared to the one calculated with more gradient-directions), it succeeds in providing structural information of the white matter with just 3 acquired directions.

We must recall that this method is not initially intended to carry out clinical studies or to obtain detailed anisotropy information, but simply to complement existing acquisition methods with an anisotropy measure. The acquisition remains unchanged, only some extra processing is needed.

## 2 METHODS

### 2.1 Diffusivity Anisotropy

In [4], authors proposed a series of advanced anisotropy measures that could be calculated from a single shell acquisition. Among them, the Diffusion Anisotropy (DiA) was presented as a robust alternative to the FA. The DiA assumes a Gaussian diffusion profile for the normalized magnitude signal provided by the MRI scanner, *E*(**q**):

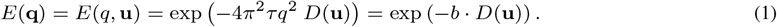

where the positive function *D*(**q**) = *D*(*q, θ, ϕ*) is the *diffusivity signal*, also known as the Apparent Diffusion Coefficient (ADC), *b* = 4*π*^2^*τ* | | **q** | | ^2^ is the b-value, *τ* is the effective diffusion time and *q* =| |**q**| | and *θ, ϕ* are the angular coordinates in the spherical system. Note that, in this case, we constraint the radial behavior so that the diffusivity *D*(**q**) does not depend on the radial direction: *D*(**q**) = *D*(**u**), where | | **u** | | = 1 and **q** = *q***u**.

Under this assumption, the DiA is defined as [4]

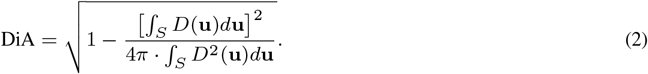

The integration on the surface of the sphere from a limited number of samples can be performed by fitting corresponding signals in the basis of Spherical Harmonics (SH), whose 0-th order coefficient is defined as:

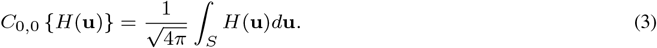

This way, a practical implementation of DiA was originally defined as

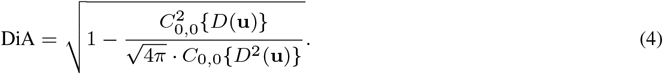

This implementation can be seen as a generalization of the Coefficient of Variation of the Diffusion (CVD) defined in [5] and an alternative definition to the Generalized Anisotropy proposed in [6].

### 2.2 Simplified DiA and Color-by-orientation

The advantage of the definition of DiA using a SH base, like the one proposed in eq. (4), is that the integral can be roughly estimated from just 3 orthogonal values. Let us assume that we only considered 3 orthogonal gradient directions in the scanner and we can extract three diffusivity values per pixel, *D*_*x*_(**x**), *D*_*y*_ (**x**) and *D*_*z*_ (**x**). We can calculate the *average diffusivity* by simply averaging the three components.

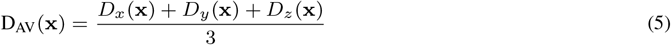

At the same time, DiA can be calculated using eq. (4):

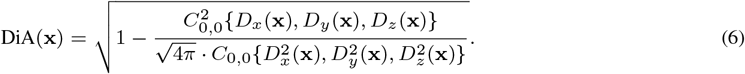

However, since the three acquired DWIs are orthogonal, the DiA can be alternatively calculated using the simplified formulation for CVD defined in [5]:

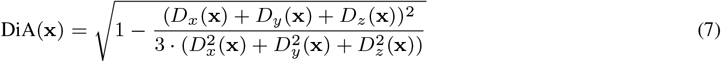

Note that, in this case, since we are assuming three orthogonal vectors, we do not need to know the value of the gradient directions in order to calculate the DiA.

We can also provide color-coded anisotropy information using DiA. In DT imaging, the anisotropy is usually coded using a RGB color system [7] in which blue is superior-inferior, red is left-right, and green is anterior-posterior. For visual purposes, the luminance of the color is weighted by the FA. Analogously, we define the RGB components as a function of the three orthogonal directions normalized by the average diffusivity, so that:

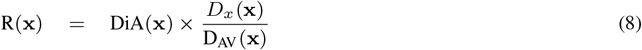

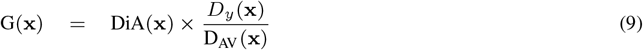

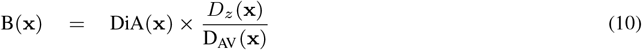

The calculation of the different metric from the 3 acquired orthogonal measures is surveyed in Fig. 1.

**FIGURE 1.**
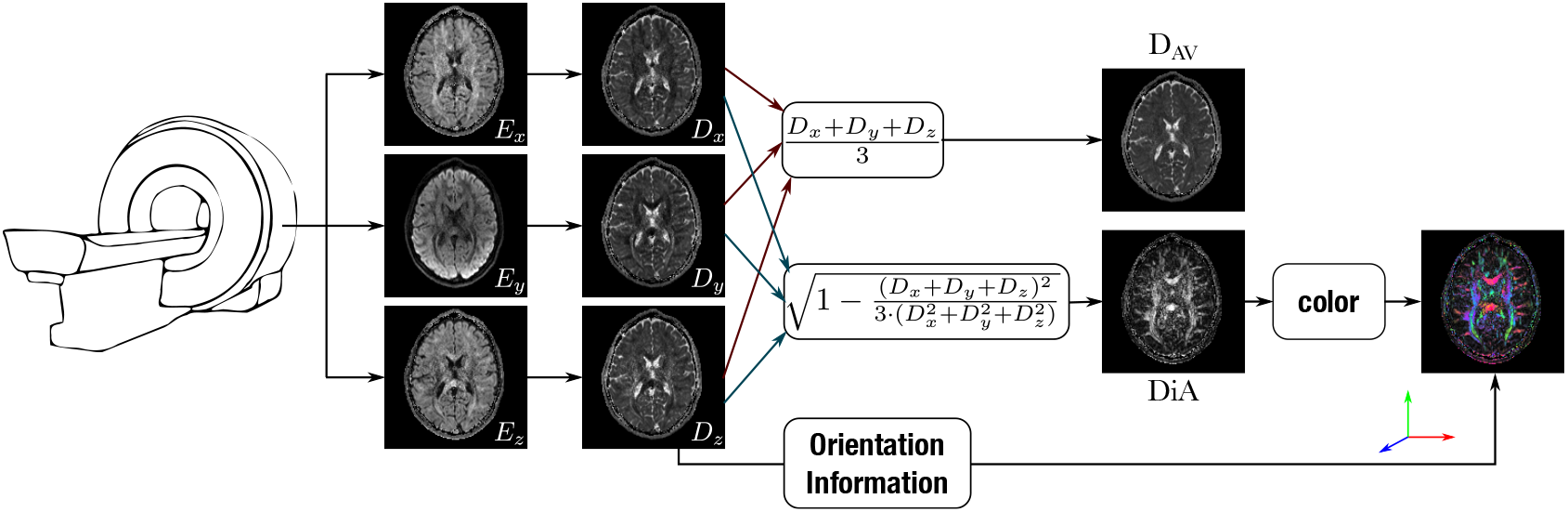
Scheme of the calculation of the different metrics derived from 3 DWIs acquired with 3 orthogonal gradient directions.

## 3 RESULTS

### 3.1 Data used for the experiments

In order to test the proposed measures, the **Human Connectome Project (HCP)**^1^ database is used, specifically volumes MGH1007, MGH1010 and MGH1016, acquired on a Siemens 3T Connectom scanner with 4 different shells at *b* =[1000, 3000, 5000, 10000] s/mm^2^, with [64, 64, 128, 256] gradient directions each, in-plane resolution 1.5 mm and slice thickness 1.5 mm. We will only make use of the innermost shell (*b* = 1000 s/mm^2^ and 64 gradient directions).

### 3.2 Visual Assessment

First, we calculate the proposed metric over 3 slices (42, 52 and 65) from the HCP volume MGH1010. The D_AV_ and DiA were calculated using only 3 DWIs from the shell at b=1000 s/mm^2^. In order to downsample to 3 gradients among the original 64, we carry out an exhaustive search of the set of three values closer to an orthogonal disposition. DiA is calculated using eq. (4); the SH are fitted with a Laplace-Beltrami penalty *λ* = 0.006. For the sake of comparison, we have also calculated the FA at b=1000 s/mm^2^ with 64 and 6 gradient directions and DiA with 64 directions. Results are shown in Fig. 2.

**FIGURE 2.**
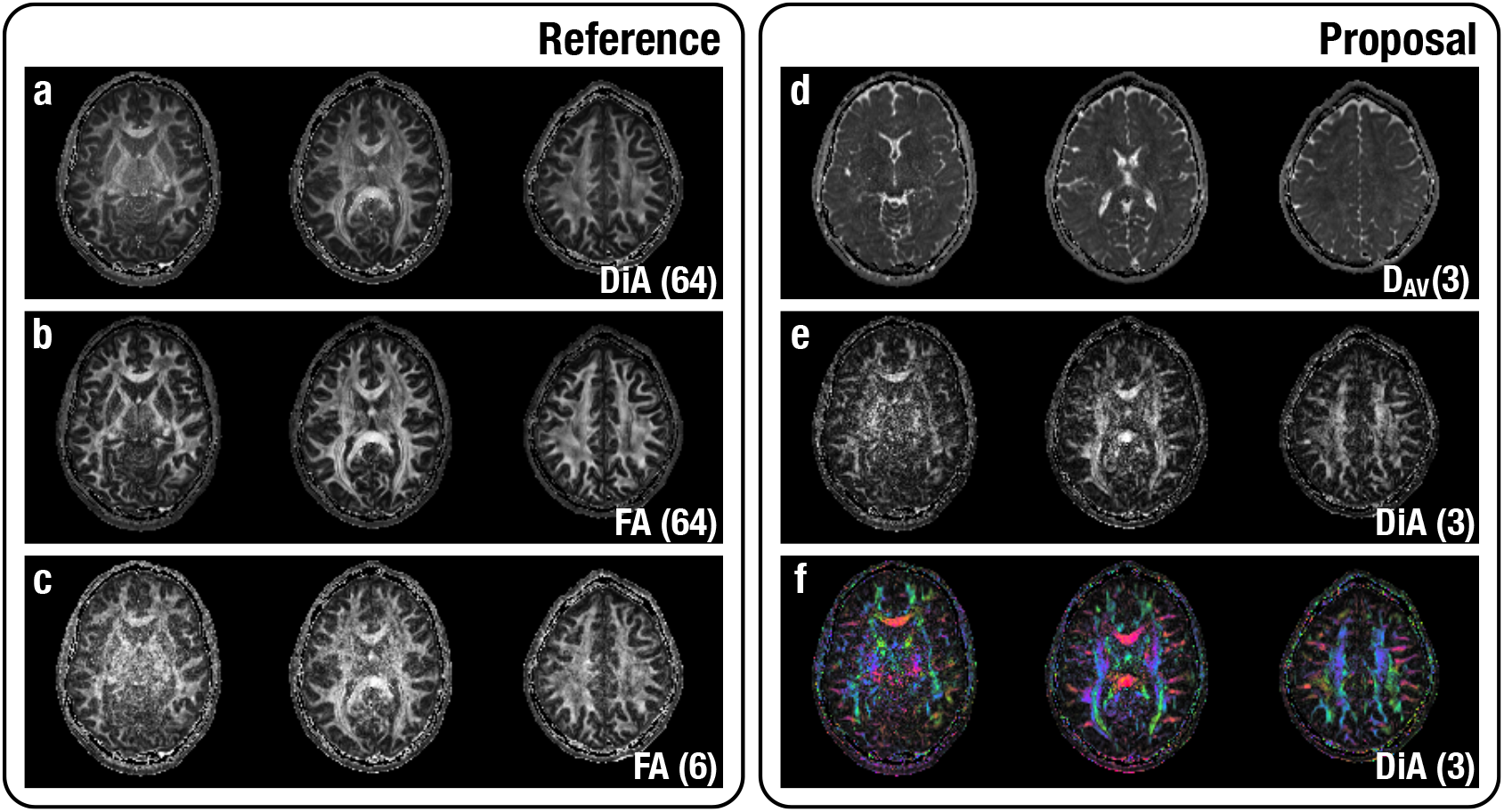
Visual assessment of proposed methods. Slices (42, 52, 65) from the HCP volume MGH1010 are shown. For the sake of comparison we have added (a) DiA (using 64 gradient directions); (b) FA (using 64 gradient directions; (c) FA (using 6 gradient directions. The proposed metrics are calculated with 3 gradient directions: (d) Average Diffusivity; (e) DiA; (f) DiA with orientation color code.

Although the visual quality of DiA calculated with 3 directions, Fig. 2-(e), is clearly poorer than FA and DiA with 64 directions (which is obvious), note that DiA succeeds in estimating information about orientation and anisotropy with just 3 gradient directions. Main structures are visible within the white matter, even clearer in the colored version, Fig. 2-(f). Thus, the same fast acquisition that can produce information about the amount of diffusion can also be used to provide rough information about the orientation of such diffusion.

Next, we compared the two ways of calculating DiA: (1) using SH and eq. (4) and (2) using the simplified approach in eq. (7). Note that the former needs the gradient directions and a SH fitting. The latter can only be used if the orientations are orthogonal and implicitly assumes that the 3 orientations correspond to the axis *x, y* and *z*. Once again we have 3 slices (42, 52, 65) from a different HCP volume, MGH1007. DiA was calculated using only 3 DWIs at b=1000 s/mm^2^ obtaining by the same downsampling we did before. Results are shown in Fig. 3, together with the absolute error between the two approaches in Fig. 3-(c). Results for both methods are totally alike. There is a small error produce by the fact that the selected directions are not totally orthogonal. Note that DiA is bounded in the range [0, 1], with an average value of 0.23 inside the brain and the average error in the same area is 1.6 *·* 10^*−*4^. In the figure we can see that the maximum error is around 1.1 *·* 10^*−*3^, which confirms that the difference between implementations is really small.

**FIGURE 3.**
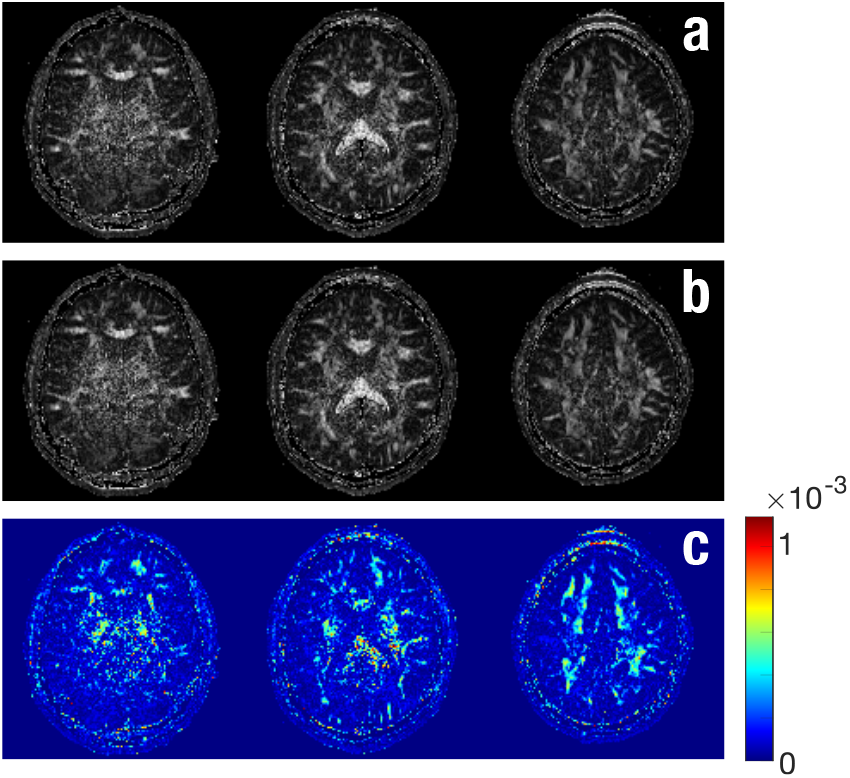
DiA using different methods on MGH1007. Both cases calculated using 3 orientations: (a) DiA using SH fitting in eq. (4); (b) DiA using the simplified expression in eq. (7); (c) Absolute error between both methods.

### 3.3 Numerical assessment

Next, we quantified the loss of information in DiA when calculated using only 3 different orientations. First, we tested the dependency of DiA on the number of diffusion samples taken in a given shell. To that end, we used a whole volume from the HCP data, MGH1016. The volume was divided in 6 different regions according to their diffusion features. The DiA was first calculated at b=1000 s/mm^2^ using 64 directions and those voxels with DiA< 0.1 removed. The remaining voxels were clustered in 6 different groups using k-means. Each voxel in the white matter was assigned to one cluster using its DiA value and the minimum distance. The following test was carried out: we began with the 64 samples (gradient directions) and uniformly downsampled this set to obtain either 3, 6, 15, 24, 35 and 48 diffusion directions subsets^2^. The DiA was computed for each considered case, and the median value inside each of the six clusters is depicted in Fig. 4. Although DiA shows it is a consistent measure when it is calculated using over 20 different orientations, for fewer gradient directions this measure is underestimated. This effect is much more noticeable when using only 3 directions. However, note that the separation between clusters remains constant. This means that the differences in the anisotropy detected by these measures can still be seen when using 3 orientations.

**FIGURE 4.**
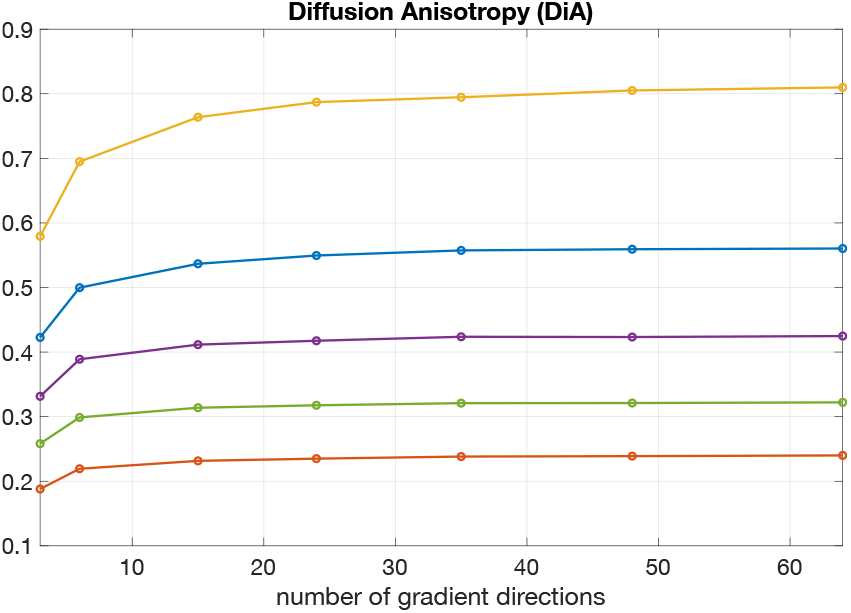
Evolution of DiA with the angular resolution (number of gradient directions), using data from HCP. The volume has been clustered in 6 different sets (for original DiA with 64 directions) and the median of each set is shown. Centroids of the data *C*_*L*_ = {0.24, 0.32, 0.42, 0.57, 0.82}.

Finally, in order to test how much information we are losing by reducing the number of gradients, we have used the same data, but 50 different random configurations were considered for each number of gradients. We have calculated two metrics:

1. The **Coefficient of Variation** (CV) for the different realizations:

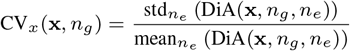

where *n*_*e*_ = 1, …, 50 is the number of experiment, *n*_*g*_ = 3, …, 64 is the number of gradient directions selected and *std* and *mean* denote the sample standard deviation and the sample mean. In order to get a single value for each direction, the median along the image is considered:

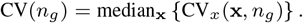

The CV is a measure of the variability of DiA for a particular number of gradients.
2. In order to quantify the difference between the metric calculated with a high number a orientations (64 in this case) and fewer directions, we have considered the **relative absolute error** (RAE) defined as:

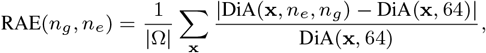

where |Ω| is the number of points considered in the volume and DiA(**x**, 64) is the DiA calculated using the 64 gradient directions. The median of all the experiments is considered:

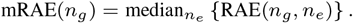

For the sake of robustness, only those voxels in which DiA(**x**, 64) > 0.2 are considered. Results are depicted in Fig. 5. As expected, when the number of gradients decrease there is an increment in the variability of the data (Fig. 5-a) and the absolute error (Fig. 5-b). It is specially significant that there is an important worsening of the image when compared to 8 gradients but we must consider that only 3 orientations have been used.

**FIGURE 5.**
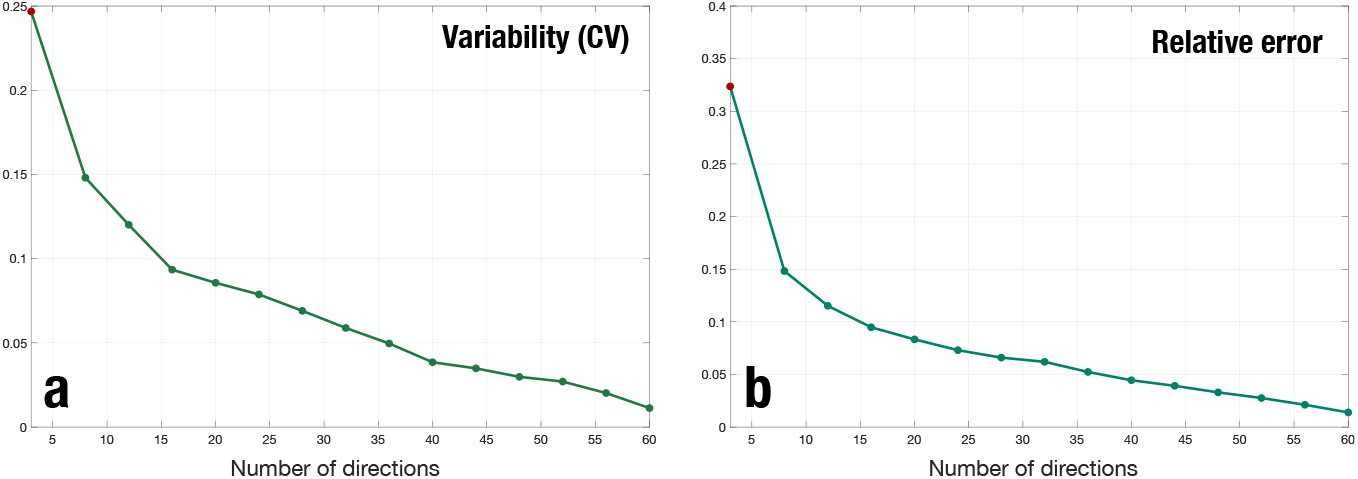
Variability and error of the DiA as a function of the number of gradient directions. In red, the value for 3 gradient directions. (a) Coefficient of variation; (b) relative absolute error.

## 4 DISCUSSION

The calculation of anisotropy measures over diffusion data is usually limited by the 6 gradient directions needed to estimate the components of the diffusion tensor. Hence, in fast acquisitions with fewer acquired orientations, only information about the *amount* of diffusion is usually provided. That is the case of fast diffusion sequences in comercial scanners in which the installed software provides an image which is the average of 3 images acquired with 3 ortogonal gradient directions where no information about the orientation of the diffusion is provided.

In this work we have proposed a method that is able to calculate a rough anisotropy image that could give information about the anisotropy and orientation of the diffusion with just 3 orthogonal directions. The method is totally compatible with existing fast acquisitions and it does not need extra data or information to be calculated. The same DWIs used for the MD estimation can provide an anisotropy metric.

The intention of the method proposed here is not to be used in clinical studies or as a substitute of the FA, but to provide complimentary information in fast diffusion acquisitions. One might argue that the DiA calculated from only 3 DWIs shows a great variability and a noisy behavior. This is by no means something new to diffusion imaging: the effect of the reduction of the number of directions has been previously studied [8, 9]. When using only 3 directions, the effect must be more noticeable. On the other hand, the main advantage of the proposed method is, precisely, the ability to provide anisotropy information with the smallest possible data set.

## 5 CONCLUSIONS

The calculation of anisotropy information from DMRI is feasible using fewer than 6 gradient directions by using alternative metric that avoids the DT calculation. In this work, we have proposed the use of DiA. This metric succeeds in providing a FA-like image from only 3 orthogonal images.

## Software

The full implementation of DiA is included in the AMURA toolbox and it may be downloaded for Matlab^*©*^ and Octave, together with use-case examples and test data, from: http://www.lpi.tel.uva.es/AMURA.

## Acknowledgements

This work was supported by Ministerio de Ciencia e Innovación of Spain with research grant RTI2018-094569-B-I00. Data collection and sharing for this project was provided by the *Human Connectome Project* (HCP; Principal Investigators: Bruce Rosen, M.D., Ph.D., Arthur W. Toga, Ph.D., Van J. Weeden, MD). HCP funding was provided by the National Institute of Dental and Craniofacial Research (NIDCR), the National Institute of Mental Health (NIMH), and the National Institute of Neurological Disorders and Stroke (NINDS). HCP data are disseminated by the Laboratory of Neuro Imaging at the University of Southern California.

## Footnotes

1 Data obtained from the Human Connectome Project (HCP) database (ida.loni.usc.edu/login.jsp) . The HCP project (Principal Investigators: Bruce Rosen, M.D., Ph.D., Martinos Center at Massachusetts General Hospital; Arthur W. Toga, Ph.D., University of Southern California, Van J. Weeden, MD, Martinos Center at Massachusetts General Hospital) is supported by the National Institute of Dental and Craniofacial Research (NIDCR), the National Institute of Mental Health (NIMH) and the National Institute of Neurological Disorders and Stroke (NINDS). HCP is the result of efforts of co-investigators from the University of Southern California, Martinos Center for Biomedical Imaging at Massachusetts General Hospital (MGH), Washington University, and the University of Minnesota.

2 A “uniform” downsampling of *n* gradients among the original 64 is here defined as those *n* directions that minimize the overall electrostatic repulsion energy among all 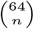 combinations. The optimization is carried out using heuristic rules.

## References

[1] de Figueiredo EH, Borgonovi A, Doring TM. Basic concepts of MR imaging, diffusion MR imaging, and diffusion tensor imaging. Magnetic resonance imaging clinics of North America 2011;19(1):1–22.

[2] Westin CF, Maier SE, Mamata H, Nabavi A, Jolesz FA, Kikinis R. Processing and visualization for diffusion tensor MRI. Medical image analysis 2002;6(2):93–108.

[3] Basser P, Pierpaoli C. Microstructural features measured using diffusion tensor imaging. J Magn Reson B 1996;111(3):209–219.

[4] Aja-Fernández S, Tristán-Vega A, Jones DK. Apparent propagator anisotropy from single-shell diffusion MRI acquisitions. Magnetic Resonance in Medicine 2021;85(5):2869–2881.

[5] Aja-Fernández S, Pieciak T, Tristán-Vega A, Vegas-Sánchez-Ferrero G, Molina V, de Luis-García R. Scalar diffusion-MRI measures invariant to acquisition parameters: a first step towards imaging biomarkers. Magn Reson Imag 2018;53:123–133.

[6] Özarslan E, Vemuri BC, Mareci TH. Generalized scalar measures for diffusion MRI using trace, variance, and entropy. Magn Reson Med 2005;53(4):866–876.

[7] Pajevic S, Pierpaoli C. Color schemes to represent the orientation of anisotropic tissues from diffusion tensor data: application to white matter fiber tract mapping in the human brain. Magnetic Resonance in Medicine: An Official Journal of the International Society for Magnetic Resonance in Medicine 1999;42(3):526–540.

[8] Barrio-Arranz G, de Luis-García R, Tristán-Vega A, Martín-Fernández M, Aja-Fernández S. Impact of MR acquisition parameters on DTI scalar indexes: a tractography based approach. PloS one 2015;10(10):e0137905.

[9] Lebel C, Benner T, Beaulieu C. Six is enough? Comparison of diffusion parameters measured using six or more diffusion-encoding gradient directions with deterministic tractography. Magnetic resonance in medicine 2012;68(2):474–483.

